# The pink salmon genome: uncovering the genomic consequences of a strict two-year life-cycle

**DOI:** 10.1101/2021.08.05.455323

**Authors:** Kris A. Christensen, Eric B. Rondeau, Dionne Sakhrani, Carlo A. Biagi, Hollie Johnson, Jay Joshi, Anne-Marie Flores, Sreeja Leelakumari, Richard Moore, Pawan K. Pandoh, Ruth E. Withler, Terry D. Beacham, Rosalind A. Leggatt, Carolyn M. Tarpey, Lisa W. Seeb, James E. Seeb, Steven J.M. Jones, Robert H. Devlin, Ben F. Koop

## Abstract

Pink salmon (*Oncorhynchus gorbuscha*) adults are the smallest of the five Pacific salmon native to the western Pacific Ocean. Pink salmon are also the most abundant of these species and account for a large proportion of the commercial value of the salmon fishery worldwide. A strict two-year life-history of most pink salmon generates temporally isolated populations that spawn either in even-years or odd-years. To uncover the influence of this genetic isolation, reference genome assemblies were generated for each year-class and whole genome re-sequencing data was collected from salmon of both year-classes. The salmon were sampled from six Canadian rivers and one Japanese river. At multiple centromeres we identified peaks of Fst between year-classes that were millions of base-pairs long. The largest Fst peak was also associated with a million base-pair chromosomal polymorphism found in the odd-year genome near a centromere. These Fst peaks may be the result of centromere drive or a combination or reduced recombination and genetic drift, and they could influence speciation. Other regions of the genome influenced by odd-year and even-year temporal isolation and tentatively under selection were mostly associated with genes related to immune function, organ development/maintenance, and behaviour.

## Introduction

Pink salmon are an economically important species under heavy exploitation and have been the subject of intense mitigation efforts. Commercial catches of pink salmon comprise roughly half of all Pacific salmon catches by weight and a much greater percentage by count as they are the smallest of the commercially important Pacific salmon (1, 2). Since the late 1980s, more than a billion pink salmon are released annually from hatcheries (1) to maintain the abundance of this fishery.

The native range of pink salmon encompasses parts of the southern Arctic Ocean between North America and Asia as well as much of the northern Pacific Ocean (3). Recently, Arctic climate warming has opened previously inaccessible Arctic territory to pink salmon as well (4–6). Pink salmon have been introduced to the Great Lakes in North America (7) and drainage basins of the White Sea (reviewed in (8)), near the border of Russia and Finland.

Pink salmon spend a year and a half at sea before returning to rivers to spawn at two-years of age. This strict two-year life history, unique to this species among salmon, has wide-ranging implications for their evolution, conservation, and possibly for their future as a species. Populations that spawn in an even-numbered year (e.g., 2020) return in an even-year (e.g., 2022) to spawn as adults. Similarly, odd-year spawned pink salmon return in odd-numbered years. Gene flow between year-classes/lineages is consequently limited (9) (this phenomenon is known as allochronic or temporal isolation). Rare exceptions to a strict two-year life-cycle of pink salmon in their native range have been reported (10–12). Outside their native range, three-year-old pink salmon have been observed in the Great Lakes following introduction (7, 13). One hypothesis based on experimental rearing in heated sea water is that temperature may play a role in precocious development (i.e., one-year life-cycle) (14).

Within a year-class, population genetic differentiation among rivers tends to be lower than that of other salmon species, which is a possible consequence of increased straying of pink salmon from natal streams during spawning (15, 16). Increased straying itself may be a repercussion of the reduced time that pink salmon spend in their natal streams compared to most other salmon species (chum salmon – *Oncorhynchus keta* being an exception) (3). Pink salmon are ready for sea migration as soon as they emerge from gravel and after yolk-sac absorption (17).

In contrast to the regional reduced heterogeneity observed within year-class populations, there is a high level of divergence between year-classes as a result of limited gene flow (9,18–22). Genetic differentiation between odd and even lineages from the same river is greater than within year-class differentiation, a phenomenon observed across the species natural range (23). There are also phenotypic differences that have been reported between lineages such as gill raker counts (19), length/size (with even-year fish tending to be smaller in Canada) (24–26), and survival/alevin growth in low temperature environments (27).

The divergence of pink salmon from other Pacific salmon species has been estimated to have occurred several million years ago (28–32); this provides a maximal time of odd and even lineage divergence. Based on mitochondrial nucleotide diversity, divergence times between odd and even-year lineages have previously been estimated as 23,600 years (33), 150 – 608 thousand years ago (34), and 0.9 – 1.1 million years ago (22). The relatively recent estimates of divergence are inconsistent with complete temporal isolation between odd and even lineages (potentially for several million years). It has been suggested that low-level gene flow or recolonization of extripated year-classes by alternate year-classes could account for recent estimates of divergence, with recolonization being a favoured explanation (33). Both low-level gene flow and recolonization have been observed in introduced pink salmon in the North American Great Lakes (7,35,36), revealing that it is possible that environment and temperature (suggested in (36)) can alter the strict allochronic isolation observed in modern times. Maturation in pink salmon has been verified to be sensitive to temperature and photoperiod under experimental conditions (37).

While odd-year and even-year pink salmon populations may occupy the same environment (during different years), these lineages can still have different selective pressures (38). For example, the density of pink salmon is known to vary between years (38, 39), and density may influence the composition of pink salmon predators, prey, and the number of fish on the spawning grounds (40–42). In years with a high abundance of pink salmon, some studies have reported a decrease in body size of pink salmon at sea (other species of salmon and seabirds have also been adversely influenced during these high abundance years) (41–45). These studies reveal that the intraspecific competition among other pink salmon and interspecific competition among other species can vary significantly between odd and even-years.

In this study, we present genome assemblies for both odd-year and even-year lineages, develop a transcriptome to annotate these assemblies, and analyze polymorphisms found between groups. We identified regions of the genome that have diverged between odd and even-year lineages with some possibly as a response to selection. We were also able to identify large Fst peaks adjacent to many centromeres and to verify one major fusion or deletion on LG15_El12.1-15.1 by combining polymorphism data with long-read sequencing of both year-classes. These regions of the genome are important aspects of pink salmon biology and provide greater insight into the evolutionary divergence of the lineages.

## Materials and Methods

### Animal care

Fisheries and Oceans Canada Pacific Region Animal Care Committee (Ex. 7.1) was the authorizing body for animal care carried out in this study. All salmon were reared, collected, or euthanized in compliance with the Canadian Council on Animal Care Guidelines.

### Genome assemblies

A mature male pink salmon was sampled from the Big Qualicum River Hatchery (NCBI BioSample: SAMN16688056) on September 19, 2019 (odd-year) by hatchery personnel and euthanized by concussion as specified in section 5.5 of the Canadian Council on Animal Care guidelines. A mature male pink salmon was also sampled from the Quinsam River Hatchery (NCBI BioSample: SAMN18987060) by hatchery personnel in the same manner on July 28, 2020 (even-year). We dissected liver, spleen, kidney, and heart tissues from the carcasses and flash-froze them on dry ice immediately and stored them at −80°C. We used a Nanobind Tissue Big DNA Kit (Circulomics) to isolate high-molecular DNA following the manufacturer’s protocol from multiple tissues. In addition, Short Read Eliminator Kits (Circulomics) were used to reduce the fraction of small DNA fragments in the DNA extractions following the kit protocol for DNA samples to be sequenced on Oxford Nanopore Technologies (ONT) platforms.

We generated sequencing libraries with the prepared DNA using a Ligation Sequencing Kit (SQK-LSK109 ONT) following the manufacturer’s protocol. The libraries were sequenced on a Spot On Flow Cell MK1 R9 with a MinION (ONT) or a PromethION (R9.4.1 flow cell). Libraries sequenced on the PromethION were size selected using magnetic beads (0.4:1 ratio). DNase flushes were performed to increase yield according to manufacturer’s instructions. We also tried to add 1% DMSO immediately before sequencing to reduce secondary structures that might block pores and reduce sequencing efficiency for one flow-cell (with a minor increase in pore occupancy, more titration will be needed to identify if there are benefits of adding DMSO). FASTQ sequence files were generated either using the Guppy Basecalling Software (version 3.4.3+f4fc735 for sequences from the MinION) with default settings or MinKNOW v3.4.6 (for sequences from the PromethION).

Short-read sequence data were generated for genome polishing for the even-year genome assembly (NCBI SRA accession: SRX10913279 – SRX10913282) and the odd-year genome assembly (NCBI SRA accession: SRX6595859 – SRX6595860). We generated the short-read data for the even-year genome by shearing 1ug of DNA (pink even-year male described above) with a COVARIS LE220 (Covaris) using the following configuration in a 96 microTUBE plate (Covaris): duty 20, pip450, cycles/burst 200, total time 90s, pulse spin in between 45s treatment. The library was then constructed using the MGIEasy PCR-Free DNA Library Set (MGI) following the manufacturer’s protocol. The library was then sequenced on an MGISEQ-200RS Sequencer (150 + 175 PE).

We generated the short-read sequence data for polishing the odd-year genome assembly for a previous assembly that was not published because the contiguity of the assembly was low. The sequences were from an odd-year haploid female produced at Fisheries and Oceans Canada using source material from the Quinsam River Hatchery (NCBI BioSample: SAMN12367892). To produce the haploid salmon, we applied UV irradiation (560 uW/cm^2^ for 176 s) to sperm from a Quinsam River male pink salmon (to destroy parental DNA) immediately before fertilizing eggs from a Quinsam River female pink salmon. Prior to sequencing, the individual was confirmed to be haploid using a panel of 11 microsatellites. The details of the library preparations and sequencing technology can be found on the NCBI website (NCBI SRA accession: SRX6595859 – SRX6595860).

We created a Hi-C library for the even-year genome assembly using the Arima-HiC 2.0 kit (Arima Genomics – manufacturer’s protocol) with liver tissue from the even-year male (NCBI SRA accession: SRR14496776). The library was then sequenced on an Illumina HiSeq X (PE150). A Hi-C library was only successfully generated for the even-year genome assembly.

After sequencing, we produced initial genome assemblies with the Flye genome assembler (version 2.7-b1587 – odd, 2.8.2-b1695 – even) (46) using ONT sequences (parameters: -g 2.4g, --asm-coverage 30). Racon (version 1.4.16) (47) was then used to find consensus sequences of the Flye assemblies (parameters: -u) after aligning the respective ONT reads to the assemblies using minimap2 (48) (version 2.13, parameters: -x map-ont). We polished the assemblies with Pilon (version 1.22) (49) using the following methods. Paired-end reads were filtered and trimmed using Trimmomatic (50) (version 0.38) (parameters for the odd-year reads: ILLUMINACLIP: TruSeq3-PE-2.fa:2:30:10 LEADING:28 TRAILING:28 SLIDINGWINDOW:4:15 MINLEN:200; parameters for the even-year reads: ILLUMINACLIP:TruSeq3-PE.fa:2:30:10:2:keepBothReads LEADING:3 TRAILING:3 MINLEN:36). The respective reads were aligned to each of the Racon-corrected assemblies using bwa (51, 52) (version 0.7.17) with the -M parameter and sorted and indexed using Samtools (53) (version 1.9) prior to polishing with Pilon (default parameters).

After the genome assemblies were polished, we identified the order and orientation of contigs/scaffolds on pseudomolecules/chromosomes for the odd-year genome using a genetic map (54) and synteny to the coho salmon genome (NCBI: GCF_002021735.2). Chromonomer (55) (version 1.10) was used to order the contigs/scaffolds using the genetic map (parameters: --disable_splitting).

Ragtag (56) (version 1.0.1) was used to order the contigs/scaffolds using synteny to the coho salmon genome (default parameters). We used a custom script (57) to compare the contig order files output by Chromonomer and Ragtag (.agp files) and manually reviewed the output for discrepancies. The manually curated order and .fasta files were submitted to the NCBI.

To order and orient contigs and scaffolds on pseudomolecules for the even-year genome, we mapped Hi-C reads to the polished assembly using scripts from Arima Genomics (58). The output alignment file was then converted to a .bed file using BEDtool bamtobed (version 2.27.1) (59) with default parameters and sorted using the Unix command ‘sort -k 4.’ After the Hi-C reads were mapped to the genome assembly, Salsa2 (60, 61) was used to further scaffold the contigs and initial scaffolds (parameters: -e GATCGATC,GANTGATC,GANTANTC,GATCANTC). After scaffolding, we mapped the remaining contigs and scaffolds onto pseudomolecules/chromosomes using the same strategy as for the odd-year genome assembly (see above) except a newer genetic map was used (62) (an odd-year genetic map was the only available) and the rainbow trout genome assembly (NCBI: GCF_013265735.2, (63)) was chosen for synteny. The proposed order and orientation was then reviewed manually using Juicebox (version 1.11.08) (64) before submission to the NCBI. The .hic and .assembly files used by Juicebox were produced using the pipeline from Phase Genomics (65). The nomenclature for the chromosomes was based on the linkage group from the genetic maps and from the Northern pike orthologous chromosomes in an attempt to standardize nomenclature across salmonids (66).

A BUSCO (Benchmarking Universal Single-Copy Orthologs) version 3.0.2 analysis (67) was used to assess assembly quality. We performed these analyses after polishing assemblies, but before mapping contigs/scaffolds onto chromosomes. The lineage dataset used in this analysis was actinopterygii_odb9 (4584 BUSCOs). The parameters used were: -m genome and -sp zebrafish.

A Circos plot was generated from the odd-year genome assembly using Circos software version 0.69-8 (68). We identified homeologous regions of the genome with SyMap version 5.0.6 (69) using a repeat-masked version of the assembly without unplaced scaffolds or contigs (default settings).

Repeats had previously been identified by NCBI and were masked by us using Unix commands. The output from SyMap was formatted and summarized using scripts from Christensen et al. (2018) (70). A histogram of repetitive sequence was generated using a python script (71). The Marey map (genetic map markers aligned to a genome) was generated using the methods from Christensen et al. (2018) (70). Centromere positions were taken from the genetic map after it was converted into a Marey map.

### Whole-genome re-sequencing

Samples were previously collected by Fisheries and Oceans Canada personnel from the following bodies of water (British Columbia unless otherwise noted): Quinsam River Hatchery, Atnarko River, Kitimat River Salmon Hatchery, Deena River, Yakoun River Hatchery, Snootli Creek Hatchery, Kushiro River (Japan) (S1 File). Samples were chosen to encompass odd-year and even-year samples from the same body of water or from nearby streams (even-year n=30, odd-year n=31).

We extracted DNA from tissues stored either in 100% ethanol or RNAlater (ThermoFisher) using the manufacturer’s protocol (72). Whole-genome sequencing libraries were produced at McGill University and Génome Québec Innovation Centre (now the Centre d’expertise et de services Génome Québec). The libraries were generated using the NxSeq AmpFREE Low DNA Library Kit and NxSeq Adaptors (Lucigen). They were then sequenced on an Illumina HiSeq X (PE150).

We identified nucleotide variants using GATK (73–75) (version 3.8). Unfiltered paired-end reads were aligned to the Racon corrected odd-year genome assembly (as other versions were unavailable at the time – available at: https://doi.org/10.6084/m9.figshare.14963721.v1) using bwa mem (parameters: -m) and the sort command from Samtools. Picard’s (76) (version 2.18.9) AddOrReplaceReadGroups was used to change read group information (with stringency set to lenient). Samtools was used to index the resulting alignment files, and the MarkDuplicates command from Picard was used to mark possible PCR duplicates (lenient validation stringency). The MarkDuplicates command was also used to merge .bam files if multiple sequencing lanes were used to sequence the sample. Read group information was changed using the Picard command ReplaceSamHeader for these samples so that the library and sample ID were the same, but other information was left the same. This was performed so that GATK would treat the sample appropriately.

HaplotypeCaller (GATK) was then used to generate .gvcf files (parameters: --genotyping_mode DISCOVERY, --emitRefConfidence GVCF) for each sample. The GenotypeGVCF command from GATK was then used to genotype the individuals in 10 Mbp intervals (see (77) for python script used to split into 10 Mbp intervals). The CatVariants command was used to merge the intervals afterwards. Variants were then hard-filtered using vcftools (78) (version 0.1.15) with the following parameters: maf 0.05, max-alleles 2, min-alleles 2, max-missing 0.9, remove-indels, and remove-filtered-all (VCF file available at: https://doi.org/10.6084/m9.figshare.14963739.v1). Additional filtering was done for some analyses, which are sensitive to linkage disequilibrium. Variants were filtered if heterozygous allele counts were not evenly represented — also known as allele balance (minor allele count < 20% of the major allele count, see (77) for python script). Variants in linkage disequilibrium were thinned using BCFtools (79) (parameters: +prune, -w 20kb, −l 0.4, and -n 2). Custom scripts, bwa mem, and Samtools index were used to map the variants to different genome assemblies (80).

### Transcriptome

To better annotate the genome assemblies, we collected a dataset of 19 tissues from a juvenile female pink salmon (NCBI Accessions: SRX6595821-SRX6595839). Euthanasia of this salmon was performed by placing the salmon in a bath of 100 mg/L tricaine methanesulfonate buffered with 200 mg/L sodium bicarbonate. Team dissection was used to quickly remove tissues and each tissue was stored in RNAlater Stabilization Solution (ThermoFisher) as recommended by the manufacturer.

We extracted RNA from the tissue stored in RNAlater Stabilization Solution using the Qiagen RNeasy kit (QIAGEN). Stranded mRNASeq libraries were generated at McGill University and Génome Québec Innovation Centre, with NEBNext dual index adapters. Libraries were then sequenced as a 1/39 fraction of a NovaSeq 6000 S4 PE150 lane at McGill University and Génome Québec Innovation Centre. These datasets were deposited to NCBI for use in gene annotation (BioProject: PRJNA556728).

### Population structure

As clustering techniques are sensitive to linkage disequilibrium, we used variants that were hard-filtered (including for allele balance) and filtered for linkage disequilibrium for all population structure analyses. A DAPC analysis (81) was used to cluster individuals in R (82) using the following packages: adegenet (83), vcfR (84), and ggplot2 (85). The number of DAPC clusters was determined using the find.clusters function and choosing the cluster count with the lowest Bayesian information criterion. Thirty principal components were retained with the dapc function. The variants used for the DAPC analysis were not yet mapped to chromosomes.

To complement the DAPC analysis, we also performed an Admixture (version 1.3.0) analysis (86) to identify clusters of individuals and quantify the admixture between the identified groups. To format the linkage disequilibrium thinned .vcf file, we used a custom Python script to rename the chromosomes to numbers (77) and PLINK (version 1.90b6.15) (87, 88) was used to generate .bed files (parameters: --chr-set 26 no-xy, --double-id). PLINK was also used to generate a principal components analysis. The Admixture software was then used to identify the optimal cluster number based on the lowest cross-validation error value. The admixture values from this analysis were plotted in R.

To examine population structure based on the mitochondrion sequence, we generated a phylogenetic tree based on full mitochondria sequences. The genome assembly included a mitochondrion sequence, and this region of the genome was subset from the variant file using vcftools. The resulting file and the SNPRelate (89) package in R was used to generate the phylogenetic tree.

The snpgdsVCF2GDS and snpgdsOpen functions were used to import the data, the snpgdsDiss function was used to calculate the individual dissimilarities for pairwise comparisons between samples, the snpgdsHCluster function was used to generate a hierarchical cluster of the dissimilarity matrix, the snpgdsCutTree function was used to determine subgroups, and the snpgdsDrawTree function was used to plot the dendrogram.

From the variants with minimal filtering and the variants after all filters had been applied, the heterozygosity ratio was separately calculated based on the number of heterozygous genotypes divided by the number of alternative homozygous genotypes (90, 91). The number of heterozygous and homozygous genotypes were counted using a python script from Christensen et al. (2020) (77). Heterozygous genotypes per kilobase pair (kbp) was calculated by dividing the heterozygous genotype counts by the genome size (2,528,518,120 bp) and then multiplied by 1000. This calculation was used on the variants with minimal filtering not yet mapped to chromosomes.

The number of shared alleles was calculated as a metric for relatedness using custom scripts for the variants with minimal filtering and which were mapped to chromosomes (92). This value is calculated by counting the number of alleles an individual has in common with another individual and is similar to previous work (93–95). The percent shared alleles was calculated in R (number of shared alleles divided by the total allele count multiplied by 100) and plotted using the reshape2 (96) and pheatmap (97) R packages.

Fst, nucleotide diversity (within populations—pi and between—dxy), and Tajima’s D were calculated and plotted using the R packages PopGenome (98), dplyr (99), tidyr (100), stringr, and qqman. In PopGenome, all metrics were calculated using a sliding window of 10 kbp and the data were visualized as a Manhattan plot using qqman. We used the populations module from Stacks version 2.54 (101) to calculate the number of private alleles, percent polymorphic variants, Fis (inbreeding coefficient), and Pi (nucleotide diversity within a population) for odd and even year class samples grouped as populations. These metrics were compared with the sample from Japan as an odd-year sample or as its own population. A comparison was also performed to see how filtering influenced these metrics.

### Genomic regions associated with population structure under selection

To identify regions of the genome associated with population structure identified in the DAPC analysis, we performed an eigenGWAS analysis (102). The format of the hard-filtered variants was converted to the appropriate format in PLINK, and the GEAR (103) software was used to run the eigenGWAS analysis (this was performed on a slightly different version of the genome assembly available on the NCBI, but only positions on chromosome 9 were minimally affected). Significance was corrected for using the genomic inflation factor to better identify markers potentially under selection rather than a result of genetic drift between populations. The genomic inflation factor corrected p-values were then plotted in R using the qqman (104) and stringr (105) packages. A Bonferroni correction was applied as a multiple test correction (alpha = 0.05). Only peaks with at least 5 SNPs within 100 kbp of each other were retained to reduce false-positives (nucleotide variants under selection are expected to be in linkage disequilibrium with surrounding variants and significant single variants not in linkage may be a consequence of spurious alignments).

### Sex determination and sdY

We utilized a genome-wide association (GWA) of phenotypic sex to identify the region of the genome associated with sex for all pink salmon (individual year-classes were checked as well). This analysis was also used to identify where the contig from the genome assembly with the sdY gene should be placed. This was confirmed with synteny from the rainbow trout Y-chromosome (NC_048593.1) and manual inspection of the Hi-C data (it was placed in the even-year genome assembly). The GWA analyses were performed using PLINK (parameters: --logistic --perm). Synteny was identified from alignments to the rainbow trout genome assembly (GCF_013265735.2, (63)) using CHROMEISTER (106) (default settings).

When manually genotyping the presence/absence of the sdY gene by visualizing alignments in IGV (107), we noticed some males had increased coverage of the sdY gene, and two haplotypes were identified (4 variants in non-coding DNA). The haplotypes were manually genotyped. To estimate the copy number of the sdY gene, we first used a python script to determine the average coverage of all hard-filtered variants (108). The average coverage of the four variants in the sdY gene was then divided by the average coverage of all variants.

## Results

### Genome assemblies

The odd-year assembly (GCA_017355495.1) had a combined length of ∼2.5 Gbp, with 20,664 contigs and a contig N50 of ∼1.8 Mbp. The even-year assembly had similar metrics, with a contig N50 of ∼1.5 Mbp, 24,235 contigs, and a length of ∼2.7 Gbp. We used a BUSCO analysis of known conserved genes to determine the completeness and quality of the genome assembly. Of the 4584 BUSCOs, 95.3% were found to be complete in the odd-year genome assembly (54.9% single-copy and 40.4% duplicated), 1.4% were fragmented, and 3.3% were missing. The even-year assembly also had 95.3% complete BUSCOs (51.5% single-copy and 43.8% duplicated), but more fragmented (1.6%) and fewer missing BUSCOs (3.1%).

The odd-year assembly had 26 linkage groups and extensive homeologous regions between chromosomes (Fig 1). The odd-year genome assembly contained similar levels of repetitive DNA and duplicated regions compared to other salmonids (Fig 1, (70,77,109)). Like other salmon species, increased sequence similarity was also observed at telomeres between duplicated chromosomal arms (Fig 1). Peaks of increased Fst between odd and even-year lineages were commonly found at putative centromere locations (Fig 1, Table 1).

**Fig 1.**
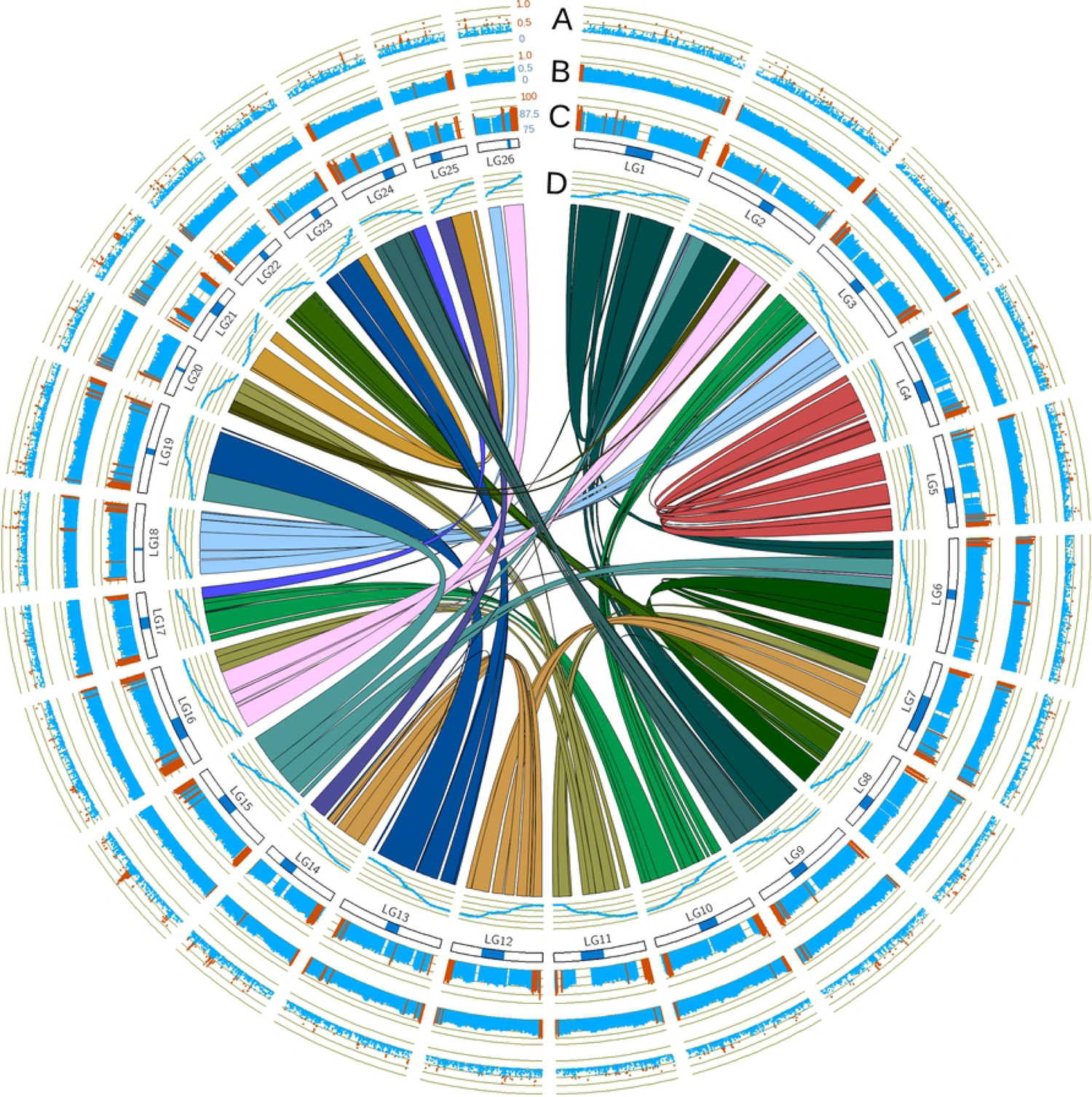
Circos plot of pink salmon genome assembly. Positions are all based on the odd-year genome assembly. Chromosomes/linkage groups are noted with blue boxes representing the centromere identified in Tarpey et al. (2017) (62). Links between chromosomes are homeologous regions identified using SyMap. A) Fst values between all odd-year and even-year salmon greater than 0.25. Values greater than 0.5 are highlighted red. B) The fraction of repetitive DNA as identified by NCBI (odd-year). Values greater than 0.65 are highlighted red. C) The percent identity between homeologous regions identified by SyMap (scale 75-100%). Values greater than 90% are highlighted red. D) A Marey map with markers from the genetic map (y-axis, 0 – 1, with 1 being the marker with the greatest cM value) placed onto the genome (x-axis, odd-year).

**Table 1.** Largest Fst peaks between odd and even-year lineages.

| Linkage group/<br>chromosome | Region<br>(Mbp) | Size of peak<br>(Mbp) | Frequency<br>Odd (p*) | Frequency<br>Even (p*) | HWE | Potential cause |
| --- | --- | --- | --- | --- | --- | --- |
| LG04_El13.1-02.1 | 50-53 | ~3 | 0.98 | 0.43 | Both | Centromere |
| LG10_El12.1-15.1 | 46.5-50 | ~3.5 | 0.69 | 0.22 | Both | Centromere |
| LG14_El18.2-23.2 | 49-55 | ~6 | 0.77 | 0.12 | Both | Centromere |
| LG15_El08.2-20.1 | 50-54 | ~4 Centromere<br>~1.26 Deletion | 0.5** | 0** | Both | Centromere/<br>Deletion-Fusion |
| LG18_El09.2-17.1 | 45.5-46.5 | ~1 | 1 | 0 | Both | Selection |
| LG21_El24.2-22.1 | 31-34.5 | ~3.5 | 0.63 | 0 | Both | Centromere |
| LG25_El23.1-24.1 | 17.5-19 | ~1.5 | 0.65 | 0.15 | Both | Centromere |
| LG26_El09.1-11.1 | 11.3-18 | ~3.5 Centromere<br>~7.3 Misassembly | 0.89 | 0.67 | Both | Centromere/<br>Misassembly |

The odd and even allele frequencies (p) were based on the most clearly defined sub-region rather than the entire region. It is unclear which individuals have the deletion or fusion on LG15\_El08.2-20.1.
\*reference genome allele frequency
\*\*alternative (to reference) allele frequency

### Population structure

A shared allele analysis (Fig 2) and both Admixture and DAPC analyses (Fig 3) revealed a clear delineation between odd and even-year lineages. Parent-progeny and sibling relationships (relationships known during sampling) are highlighted by increased levels of shared alleles, but the majority of clustering appears to be related to geographical distance (Fig 2, S1 File). No apparent admixture was observed in the even-year class (Fig 3B). In the odd-year lineage, estimated ancestry from the even-year group varied from zero to over forty percent (Fig 3B).

**Fig 2.**
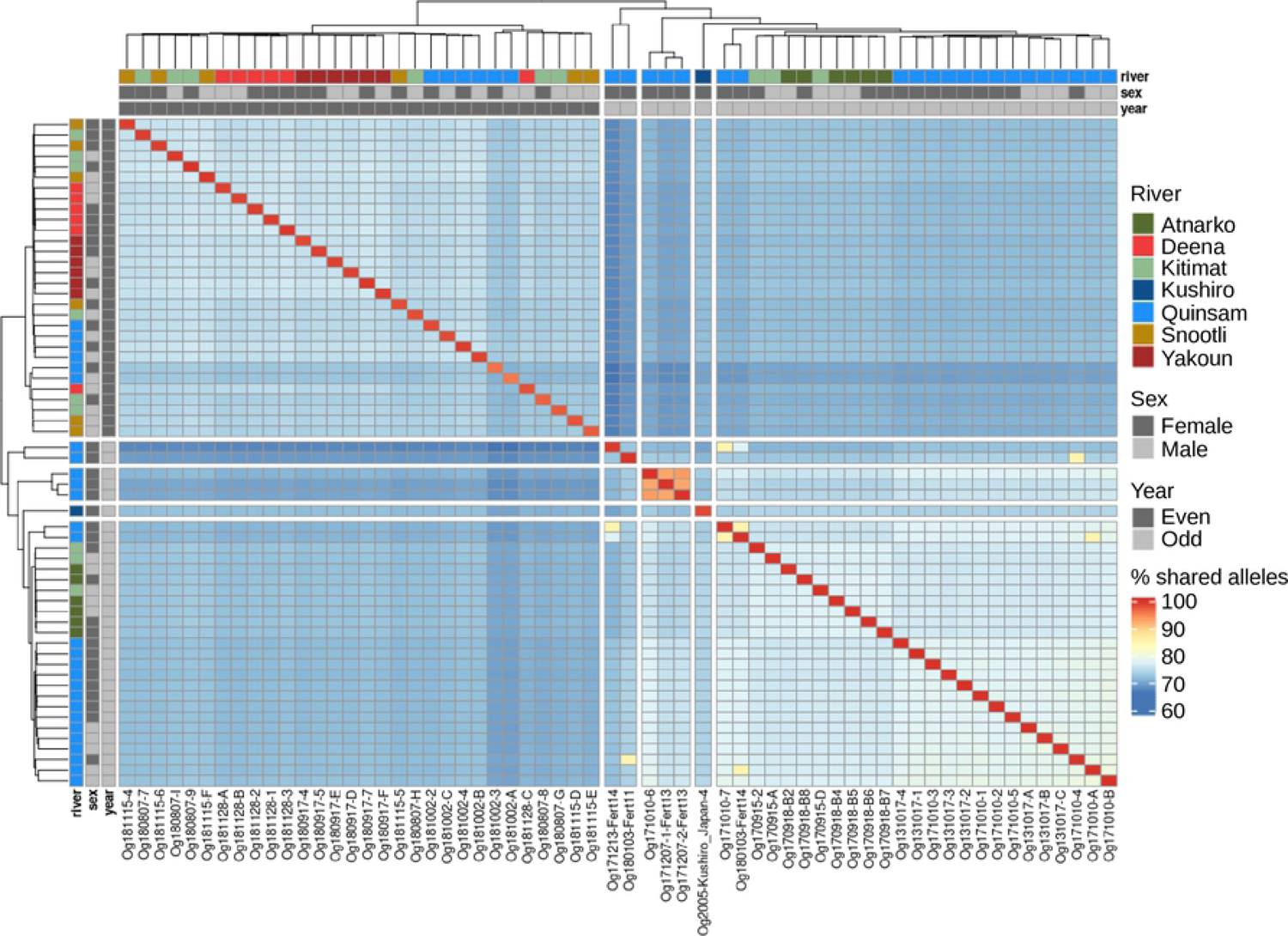
Percent of shared alleles among pink salmon. A heatmap of shared alleles between salmon is shown with clustering and a dendrogram. Each square represents the percent shared alleles after minor filtering of variants (bi-allelic SNPs). In addition to the legend displaying the colour representation of percent shared alleles, the sex, year-class, and river system sample information is colour-coded and shown on both rows and columns.

**Fig 3.**
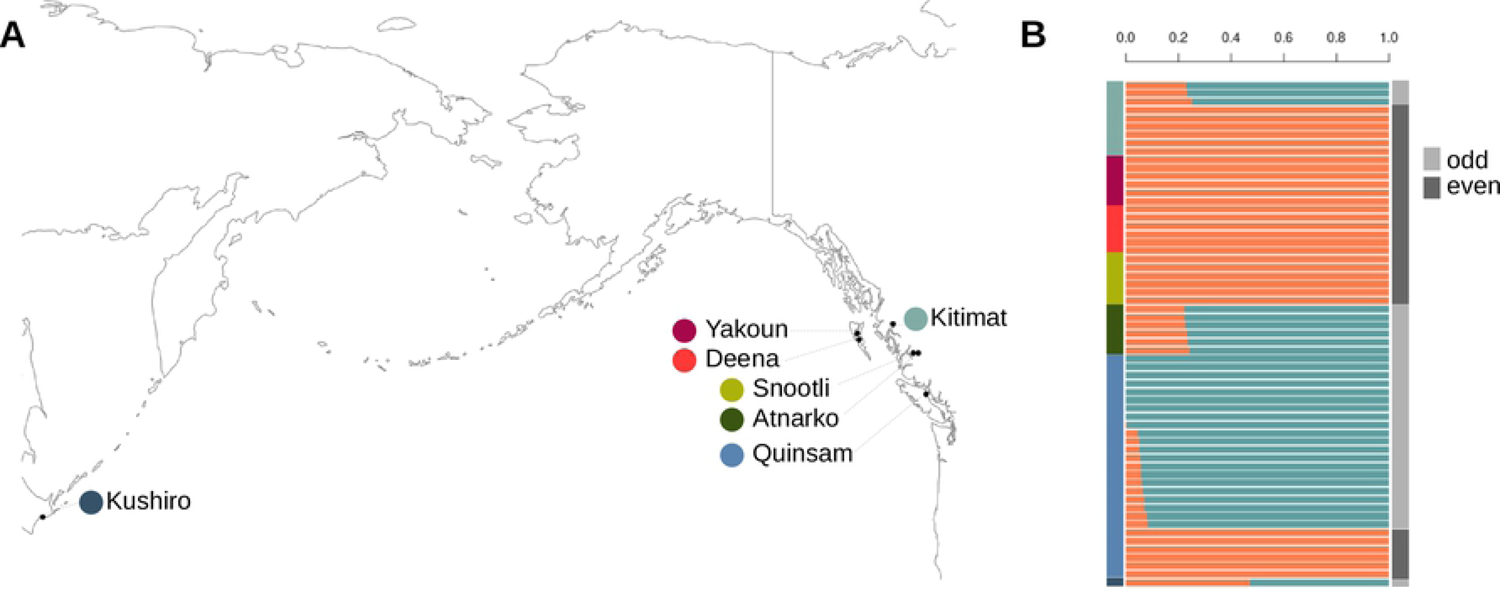
Population structure of pink salmon. A) Sampling locations for odd and even-year pink salmon. B) An admixture analysis based on an optimal group number of two. Sampling site is specified on the left (y-axis) by colour and fraction of alleles inherited from a lineage is shown on the x-axis (orange – even-year, blue – odd-year). On the right, DAPC groups are shown (see S1 File for group and coordinate positions). The DAPC groups matched year-class/lineage designations.

A separate analysis of mitochondrial DNA was performed to further investigate the relationships between the odd and even-year lineages. Odd-year pink salmon had longer branch lengths in mitochondria dendrograms and haplotype networks with more uniform distributions of haplotypes (Figs 4A and B). The even-year salmon had two major haplotypes (Fig 4B). Mitochondrial sequence analyses revealed 21 unique haplotypes from the 61 mitochondria sequences with 1-19 steps between haplotypes (Fig 4). Based on the length of the sequence analyzed (16,822 bp) this represents a mutation frequency between 0.006% to 0.1%. One haplotype was shared between lineages and the closest haplotype that was not shared had 5 steps between year-classes (Fig 4). The mitochondrial analyses illustrate divergence between the odd and even-year lineages, but also raises questions regarding possible recent admixture based on a shared haplotype and an odd-year haplotype most closely related to an even-year haplotype.

**Fig 4.**
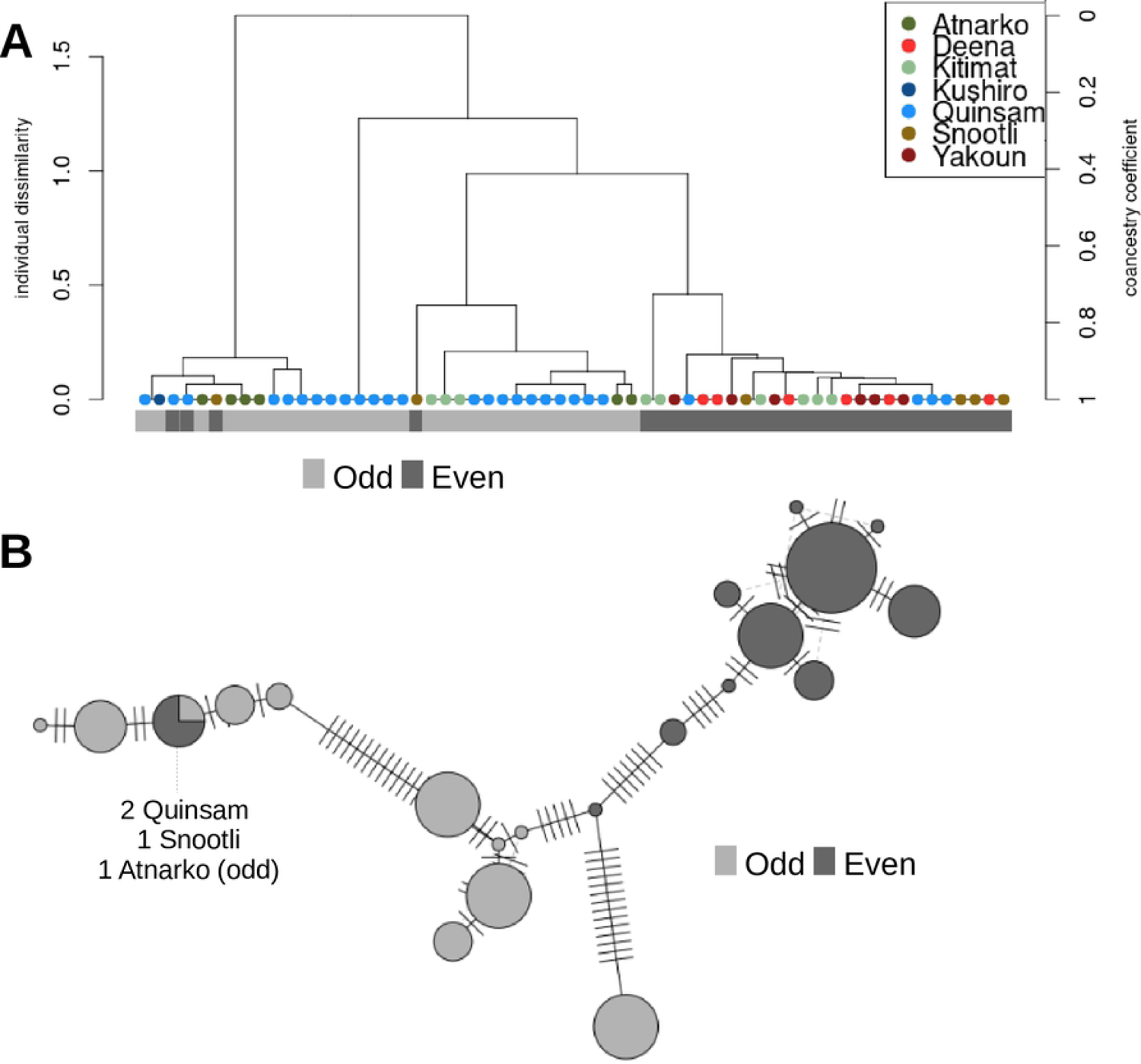
Whole mitochondrial genome comparisons between lineages. A) A dendrogram based on full mitochondrial sequences. The y-axes show dissimilarity scores on the left and coancestry values on the right, which were used to cluster individuals. Year-class/lineage is specified below the dendrogram. B) A full mitochondrial genome haplotype network is shown for the 21 unique haplotypes identified. River names are shown for the haplotype shared between lineages.

Several metrics were calculated to quantify genetic divergence between and within year-classes: heterozygosity ratios, heterozygous genotype per kbp, polymorphic sites, private alleles, and nucleotide diversity. Heterozygosity ratios in odd-year fish ranged from 1.5-4.56, with an average of 2.54 (excluding haploid individuals generated for a previous project) (S1 File). Even-year class individuals ranged from 1.09-1.78, with an average of 1.44 (S1 File). The average heterozygous genotype per kbp (excluding haploids) was 0.71 for odd-year salmon (range: 0.55 – 0.85) and 0.58 for even-year (range: 0.45 – 0.69) pink salmon. The Pearson correlation between heterozygosity ratio and heterozygosity per kb was 0.91 (excluding haploids). Salmon from odd-years had on average higher levels of polymorphic sites, increased private alleles, and increased nucleotide diversity (Table 2). These values varied based on sample inclusion and filtering parameters used for filtering nucleotide variants (Table 2). The average percent of shared alleles among odd-year fish was 76.13%, 74.42% among even-year individuals, and 71.04% between year-classes (S1 File). Most analyses revealed increased genetic diversity among odd-year pink salmon than among even-year pink salmon and fewer shared alleles between odd and even-year populations than within year-class.

**Table 2.** Population metrics of the two lineages.

|  | Odd<br>with sample from Japan | Odd<br>without sample from Japan | Even |
| --- | --- | --- | --- |
| Number of variants |  | 3,817,721 (101,594) |  |
| % polymorphic | 93.23 (97.91) | 92.61 (97.64) | 90.66 (87.86) |
| Private alleles | 356,634 (12,333) | 302,507 (10,486) | 258,335 (2,124*) |
| Pi | 0.276 (0.337) | 0.276 (0.337) | 0.269 (0.283) |
| Fis | 0.075 (0.143) | 0.068 (0.135) | 0.122 (0.195) |
| Variants with minimal filtering are shown first and variants after all filters are shown in parentheses. |  |  |  |
| *2,126 without sample from Japan |  |  |  |

### Genomic regions associated with odd and even-year lineages

We identified regions of the genome with divergence between odd and even-year lineages using an eigenGWAS and Fst analysis. Seventeen significant regions of the genome were discovered to contribute to the divergence between odd and even-year lineages (Fig 5, Table 3). These regions are putatively under selection as genetic drift is partially accounted for through the genomic inflation factor. Multiple candidate genes under selection were identified in these regions (Table 3).

**Fig 5.**
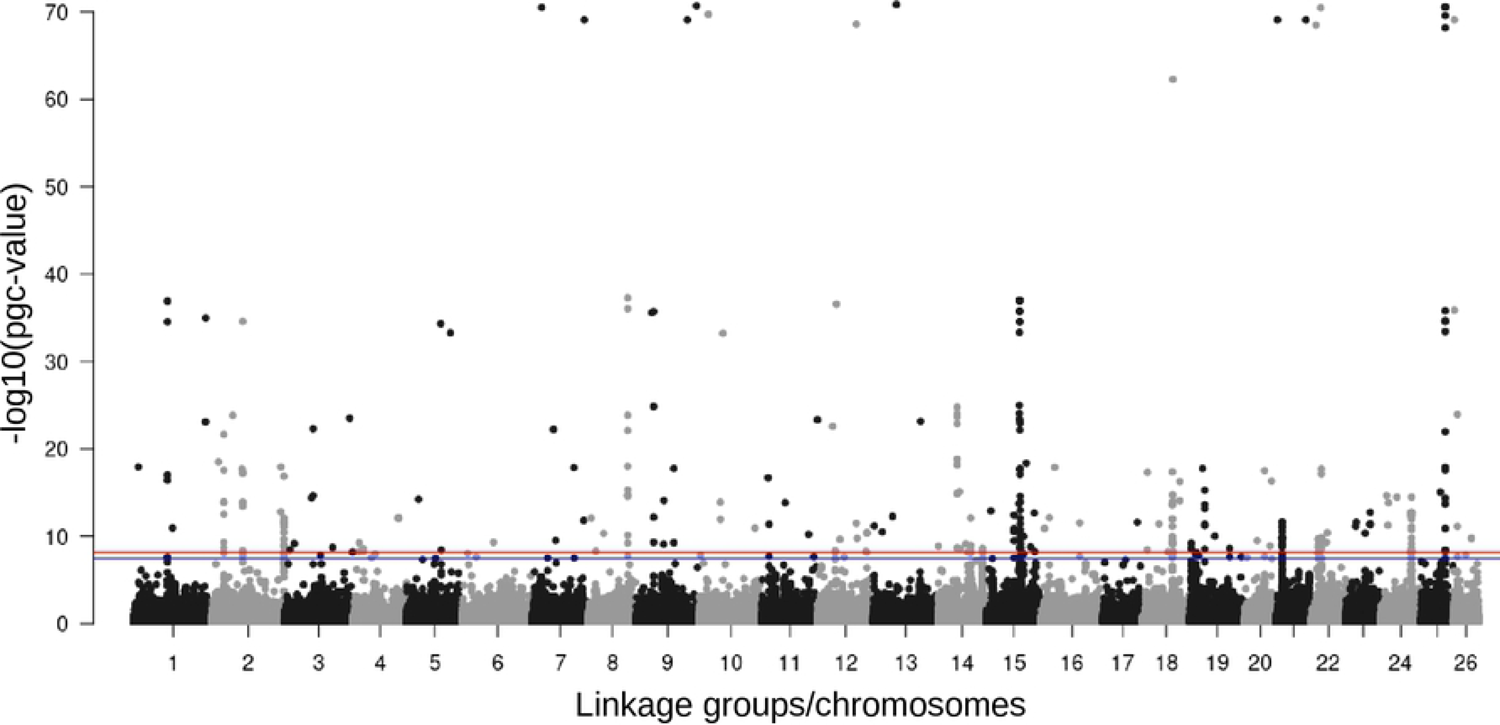
Genome-wide divergence between odd and even-year pink salmon lineages. A Manhattan plot of eigenGWAS results, with chromosome positions on the x-axis and p-values (corrected for genetic drift using the genomic inflation factor) on the y-axis identifies region of the genome potentially under selection. The red horizontal line represents a Bonferroni correction at ɑ=0.01 and the blue line at ɑ=0.05. All positions are from the odd-year genome assembly.

**Table 3.** Top eigenGWAS peaks identified between lineages.

| Chromosome | BP range | SNP position<br>with<br>lowest P-value | Candidate Gene | Gene<br>Symbol |
| --- | --- | --- | --- | --- |
| LG01_El19.1-16.1 | 51653225-51738345 | 51716026 | protein tyrosine phosphatase<br>receptor type J | *PTPRJ* |
| LG02_El19.2-07.2 | 18075760-18095551 | 18075760 | AT-rich interactive domain-containing protein 3A | *arid3a* |
| LG02_El19.2-07.2 | 46961740-47008290 | 46969254 | protein-methionine sulfoxide<br>oxidase mical2b | *mical2b* |
| LG02_El19.2-07.2 | 110392052-110493632 | 110449484 | multidrug and toxin extrusion<br>protein 2-like | *SLC47A2* |
| LG08_El03.2-06.2 | 60715584-60782108 | 60767399 | polh polymerase (DNA<br>directed), eta | *POLH* |
| LG14_El18.2-23.2 | 29365137-29414547 | 29411435 | uncharacterized gene |  |
| LG14_El18.2-23.2 | 50053735-50217619 | 50217619 | Unknown |  |
| LG15_El08.2-20.1 | 42753852-42762230 | 42758791 | cystathionine gamma-lyase | *CTH* |
| LG15_El08.2-20.1 | 51106314-52224901 | 51106314 | multiple candidates* |  |
| LG18_El09.2-17.1 | 45516859-45534019 | 45524530 | cell division control protein 42<br>homolog* | *CDC42* |
| LG18_El09.2-17.1 | 46342891-46450909 | 46347678 | H-2 class II histocompatibility<br>antigen, A-U alpha chain* | *H2-Aa* |
| LG19_El20.2-01.2 | 22831059-22843129 | 22836628 | B-cell receptor CD22 | *CD22* |
| LG21_El24.2-22.1 | 9281799-9398462 | 9391795 | histidine N-acetyltransferase | *hisat* |
| LG22_El03.1 | 10576168-10595358 | 10595038 | purine nucleoside<br>phosphorylase | *PNP* |
| LG22_El03.1 | 15334053-15411821 | 15405819 | multiple candidates |  |
| LG24_El10.1-25.1 | 47355926-47430898 | 47398112 | microtubule-associated protein<br>9 | *map9* |
| LG25_El23.1-24.1 | 37126251-37202999 | 37137812 | uncharacterized gene (ncRNA) |  |

All positions are relative to the odd-year genome.
\*associated with an Fst peak

Seven of the eight largest Fst peaks between year-classes were located in the vicinity of a centromere (Fig 1, Table 1). More detail is presented on one of the largest Fst peaks. This peak is also associated with a large deletion or fusion. The Fst peak on LG15_El12.1-15.1 (Fig 6A) is in Hardy-Weinberg equilibrium in the odd-year lineage (*p*=0.984 with a chi-square test), but fixed in the even-year lineage (Figs 6B and 6C). When Oxford Nanopore reads from the two year-classes were aligned to the genome assembly, a heterozygous deletion or fusion from 51,670,144 – 52,926,328 was found in this region of the odd-year male used for genome assembly (S1 Fig.). The ∼1.2 Mbp deletion/fusion may explain why the LG15_El12.1-15.1 Fst peak was one of the largest and widest (Fig 1, S1 Fig.).

**Fig 6.**
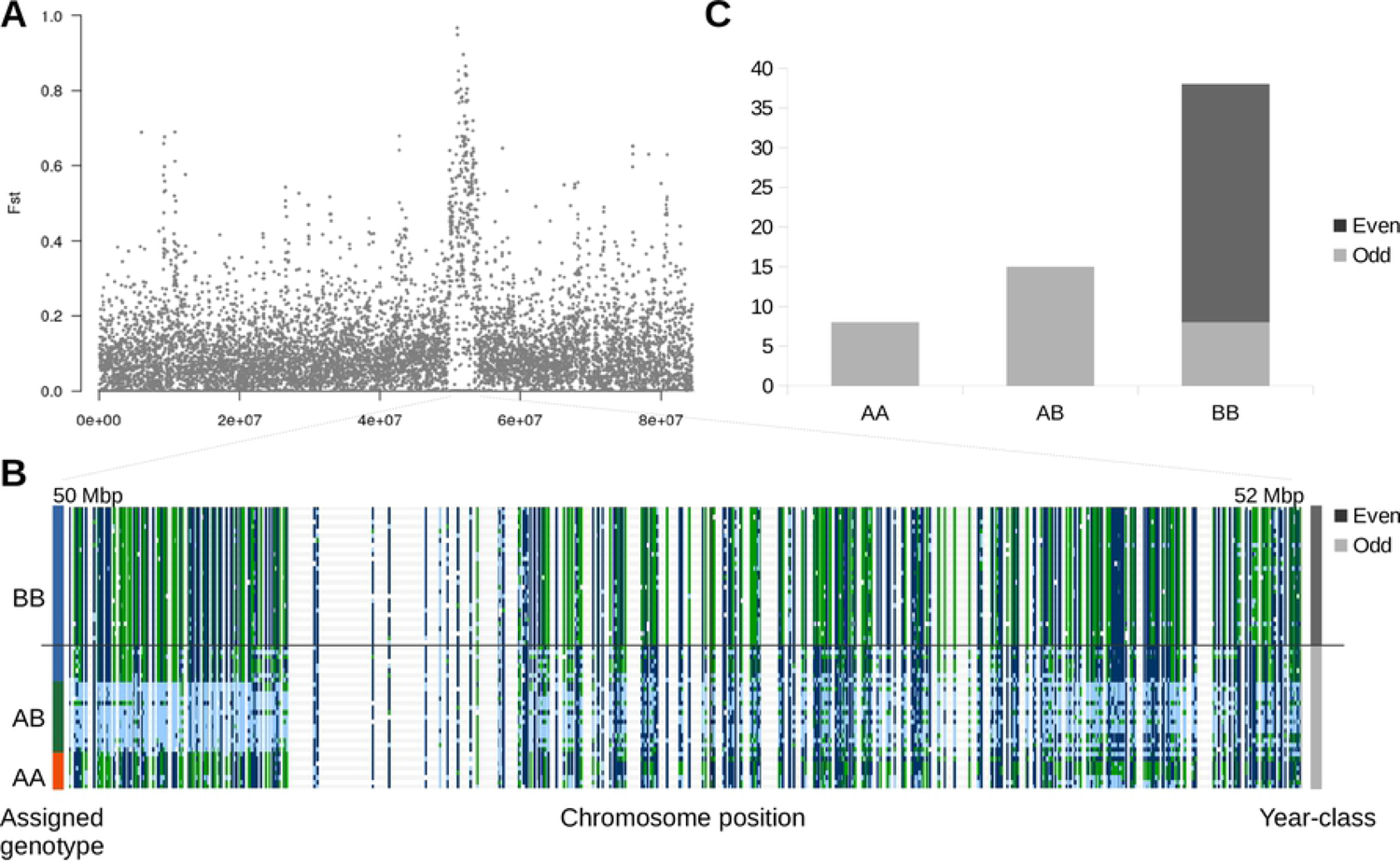
Chromosome LG15_El12.1-15.1 Fst peak. A) A Manhattan plot of 10 kbp sliding-window Fst values between odd and even-year pink salmon lineages on chromosome LG15_El12.1-15.1. B) Genotypes visualized in IGV. Each row represents an individual pink salmon and each column represents a nucleotide variant (dark blue – homozygous reference, light blue – heterozygous reference, green – homozygous alternative, and white – missing genotype). Individuals were sorted by year-class (shown on the right) and then by assigned genotype (shown on the left). C) Counts of genotypes of the chromosomal polymorphism based on manual genotyping.

It is difficult to distinguish between a deletion and a chromosomal fusion in these analyses and this may represent a fusion instead of a deletion. Previous research support chromosomal variants in pink salmon (110) and a species specific fusion of this chromosome (66), but further research will be needed to confirm this hypothesis. From these analyses, many highly divergent regions of the genome were identified, either from selection or from genetic isolation/population dynamics. The largest reservoirs of divergence between odd and even-year classes appears to be associated with centromeres, but not exclusively and uncommonly for regions putatively under selection (Table 3).

### Sex determination and sdY

The sex-determination gene in salmonids, *sdY* (111), was located on a ∼110 kbp contig in the pink salmon odd-year genome assembly (NCBI accession: JADWMN010014055.1) and on a contig ∼367 kbp that was placed onto a chromosome in the even-year genome assembly. The *sdY* gene can be placed at one of the ends of LG20_El14.2 by using genome-wide association with sex as the trait of interest, Hi-C contact data (even-year genome), and synteny with the rainbow trout Y-chromosome and chromosome 29 (an autosome) of the coho salmon (Fig 7A, S1 and S3 Figs.). LG20_El14.2, has the reverse orientation in the odd-year assembly compared to the genetic map (Fig 1), but was corrected to have the same orientation in the even-year assembly.

**Fig 7.**
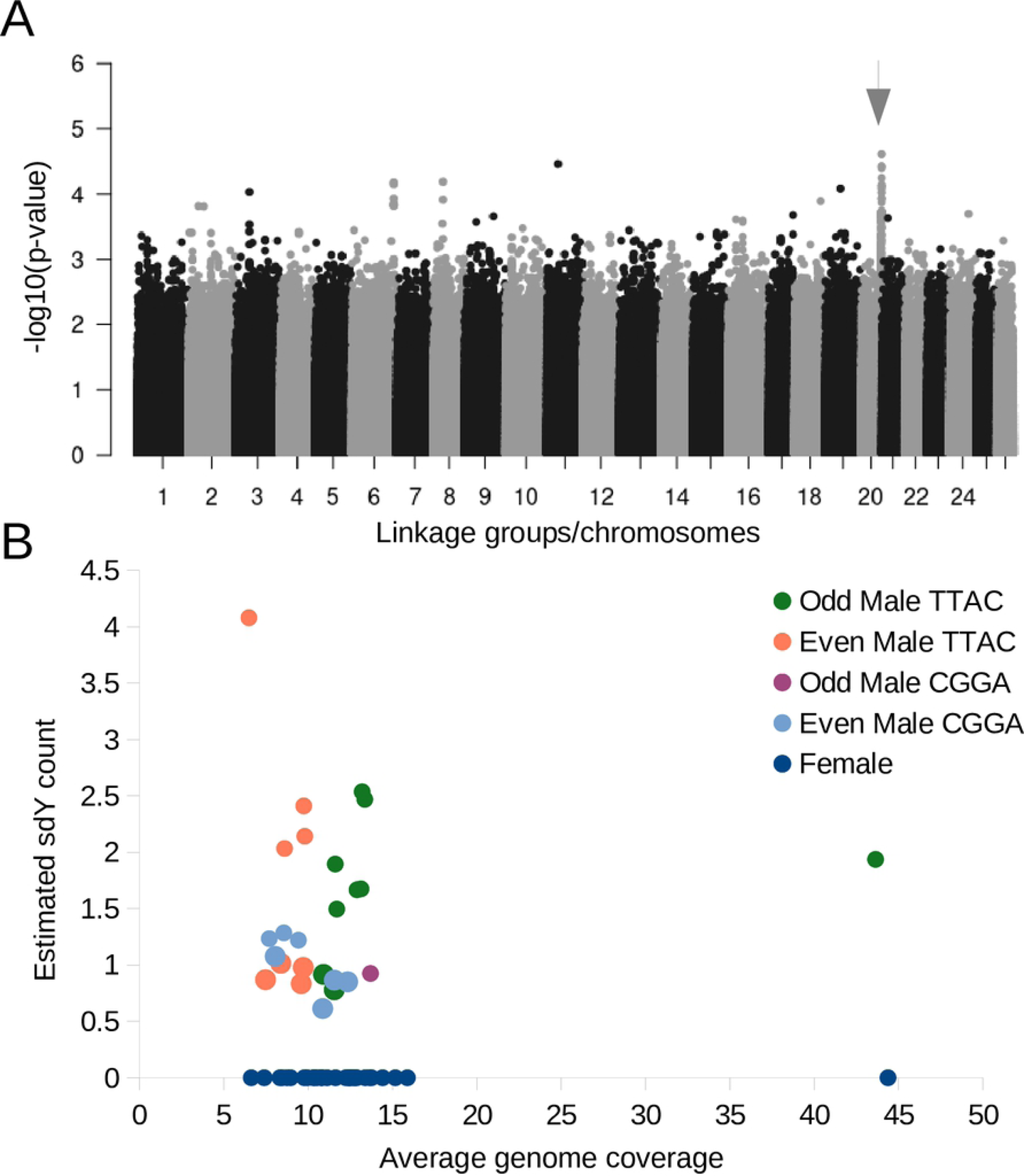
The location and counts of the sex-determining gene, *sdY*, in pink salmon. A) A genome-wide association analysis with sex as the phenotype under investigation shown as a Manhattan plot. The putative sex-determining region is indicated with an arrow. B) A scatterplot with the average coverage of the variants across the genome on the x-axis for all the pink salmon, and the estimated *sdY* count on the y-axis (*sdY* has previously been identified as the sex-determining gene in most salmonids). The different colour points represent different year-classes and *sdY* haplotypes.

In both genome assemblies there is only one copy of the *sdY* gene, confirmed with a BLAST alignment of a sdY gene available in the NCBI database (KU556848.1) to the respective assemblies. From a self-alignment of the *sdY*-containing contig, the majority of this contig is highly repetitive, > 90 kbp out of ∼110 kbp. From the alignment of the *sdY*-containing region in pink salmon to the coho salmon chromosome 29, only a small portion of the Y-chromosome appears to be unique to the Y-chromosome (S3 Fig.). Genotypes were called for the majority of this region for males and females, and the main difference related to sex was that all females had large runs of homozygosity while many males had large runs of heterozygosity (S4 Fig, S1 File).

From previous research (112, 113), a pseudo growth hormone 2 gene was shown to be tightly linked to sex-determination in pink salmon. Four tandem duplicates of this gene (NCBI: DQ460711.1) were identified on the same contig in the even-year genome assembly, but only two copies were found in the odd-year genome assembly on separate contigs (S1 File). As these contigs were not mapped to a chromosomal position, it is likely that parts of the Y-chromosome specific region remain incomplete in these two assemblies.

There were two *sdY* haplotypes (variants found in non-coding DNA) observed in both odd and even male pink salmon (Fig 7B, S1 File). Additionally, some males possessed multiple copies of the sdY gene (Fig 7B). The CGGA *sdY* haplotype was only identified in a single odd-year male pink salmon, while the TTAC haplotype was evenly distributed between year-classes (S1 File).

Based on manual inspection of the genotypes, long stretches of heterozygosity were observed near the *sdY* gene in some males, but not in others. In males with the TTAC *sdY* haplotype, there were extended or short runs of heterozygosity evenly distributed between year-classes (S1 File). Even-year males with the TTAC *sdY* haplotype and a short run of heterozygosity were more likely to have multiple copies of the *sdY* gene (n=4, average=2.7) than the same group with long runs of heterozygosity (n=4, average=0.9, p=0.017 with one-tailed, unpaired t-test). Any individuals with the CGGA *sdY* haplotype did not have stretches of heterozygosity near the putative location of *sdY*. One hypothesis to explain these results is that individuals with the CGGA *sdY* haplotype have an alternative sex chromosome.

## Discussion

### Population structure

Similar to previous studies (23, 54), pink salmon population structure divergence was found at the whole-genome level to be greater between year-classes rather than based on geography. Shared allele, DAPC, and Admixture analyses point to a clear delineation of odd and even lineages, with the exception of the only sample from Japan. Further sampling will need to be performed to provide an improved picture of the diversity of this species at the whole-genome level within year-class and across their Pacific Rim distribution. In British Columbia, however, the even-year lineage appeared to be more homogeneous than the odd-year lineage based on the admixture analysis and several population metrics such as nucleotide diversity. A similar result was previously observed with microsatellite (15) and SNP markers (23).

Divergence between lineages was also revealed by whole mitochondrial sequences. There were 21 unique mitochondria genotypes among the 61 individuals sampled, and only one of these haplotypes was shared between lineages. While the number of unique haplotypes was the same between lineages, most of the even-year class haplotypes (8 out of 10) were similar in sequence. The two major haplotypes seen in the even-year class were consistent with the Alaskan A and AA haplotypes seen in Churikov and Gharrett (2002) (33), as were the numerous and more distantly related odd-year haplotypes.

The low nucleotide diversity of mitochondrial haplotype networks and the increase of rare haplotypes have led previous studies to conclude that pink salmon (with some local exceptions) have undergone a bottleneck during the Pleistocene interglacial period and rapid expansion since the last glacial maximum or earlier (33,34,114). The interconnected mitochondrial networks in these studies have inner shared haplotypes between year-classes. Churikov and Gharrett (2002) suggested that these observations supported a model where a year-class might go extinct and an alternate year-class would then replace that population rather than continued gene flow between year-classes that would be necessary to otherwise explain the shared haplotypes (incomplete lineage sorting was tested) (33). The mitochondrial network seen in this study is consistent with that hypothesis. An alternative hypothesis is that environmental factors influence maturation timing and the strict two year life-cycle of pink salmon, and gene flow between year-classes only occurs when environmental conditions favour changes to the two year life-cycle, as that seen in the introduction of pink salmon to the Great Lakes (7,35,36).

Estimates of divergence based on mitochondrial sequences suggest that odd and even-year lineages (from East Asia and Alaska) are relatively recent for pink salmon as a species (generally less than 1 million years ago) and divergence likely began during the Pleistocene interglacial period or later (22,33,34). If the two-year life-cycle is environmentally influenced, these estimates could be distorted by phases of gene-flow and would suggest that the interglaciation period was the last major period of gene-flow between odd and even-year classes (but does not necessarily mean that was when the strict two year life-cycle evolved).

It has previously been reported that the odd-year lineage of pink salmon has higher levels of heterozygosity, private alleles, and allelic richness (23, 54). A similar trend was observed in this study with the heterozygosity ratio, heterozygous genotypes per kbp, private alleles, and other metrics assessing nucleotide diversity. Several factors could help explain the reduced levels of nucleotide diversity seen in the sampled even-year populations. Tarpey et al. (2018) suggested three possibilities, 1) the odd-year lineage was older and the even lineage was derived from the odd-year lineage, 2) there was a past reduction in even-year lineage(s), and 3) genetic variation was lost during adaptation (23).

Further sampling will be required to understand if this phenomenon is seen in all even-year populations (especially as lower heterozygosity in the even-year lineage is not universally supported, e.g. (20)).

This information is important to interpret which hypothesis is better supported or if another model is better suited (e.g., extirpated lineage replaced by alternate year or that recent population demographics are more important).

### Fst peaks between odd and even-year lineages

A single major chromosomal polymorphism (either a fusion or deletion) was identified proximal to a centromere on LG15_El12.1-15.1. This region was characterized by ∼4 Mbp runs of homozygosity/heterozygosity. This region was identified from an Fst analysis because nearly the entire region was fixed in the even-year lineage, but appeared to segregate as a single locus in Hardy-Weinberg equilibrium in the odd-year lineage.

Interestingly, pink salmon runs of homozygosity/heterozygosity were common at centromeres rather than an effect of chromosomal polymorphisms. Six other major runs of homozygosity/heterozygosity were also located near centromeres and they differed between lineages. All of these Fst peaks extend for at least 1 Mbp. It is expected that regions with reduced recombination, such as centromeres, will have increased runs of homozygosity and reduced genetic diversity (reviewed in (115)). This may help explain why there are long runs of homozygosity at centromeres, but not why there are differences between lineages at these loci. Genetic drift or selection such as centromere drive would also need to be considered.

The centromere drive hypothesis posits that a centromere can be retained in a female gamete more often than an alternative centromere during meiosis due to an advantageous DNA sequence mutation at the centromere or from mutations in centromere associated proteins (reviewed in (116–118)). In populations that become isolated, the competition between centromere sequences can quickly drive differentiation at these regions between the populations and result in hybrid defects should they come into contact again (117). These observations reveal that the pink salmon lineages may be at a point where speciation is a likely outcome as these large centromere differences could cause hybrid defects. In medaka, genomic diversity at non-acrocentric repeats in centromeres were associated with speciation (119).

The centromere drive hypothesis may further shed light on the fixation of the Fst peak on LG15_ El12.1-15.1. Robertsonian fusions (assuming that the Fst peak on LG15_El12.1-15.1 is indeed associated with a fusion rather than a deletion) can generate centromeres that are preferentially able to segregate to the egg during female meiosis (118). This could help drive the fusion to fixation in a population. Alternatively, if the telocentric chromosomes instead of the fused metacentric chromosome had more effective centromeres, the telocentric chromosomes would become fixed. Further studies will be needed to confirm if there is indeed a fusion instead of a deletion and that the fusion leads to fixation by centromere drive.

### Genomic regions putatively under selection

A large component of the genetic and phenotypic diversity between pink salmon year-classes likely originates from genetic drift as there is little evidence for gene flow between lineages. However, in addition to genetic drift, these lineages may experience different selective pressures even if they occupy the same streams. As mentioned in the Introduction, population density between lineages is often different and this can generate different ecological environments. EigenGWAS and Fst analyses were used to identify regions of the genome potentially responsive to these environmental differences between pink salmon year-classes. Candidate genes under selection were organized into three broad categories (immune system, organ development/maintenance, and behavior), and each is discussed below.

### Immune system

Variation in immune related genes is a common phenomenon between salmonid populations (e.g., (77,120,121)). Between odd and even-year pink salmon, five eigenGWAS peaks were identified near or in genes with immune related functions. These include the H-2 class II histocompatibility antigen, A-U alpha chain (*H2-Aa*) (122–126), B-cell receptor CD22 (*CD22*) (127, 128), polh polymerase (DNA directed), eta (*POLH*) (129–133), AT-rich interactive domain-containing protein 3A (*arid3a*) (134, 135), and purine nucleoside phosphorylase (*PNP*) (136–140).

Several factors could influence why these immune related genes might be under selection between odd-year and even-year populations of pink salmon. For example, altered migration patterns (reviewed in (141, 142)), increased pathogen loads between year classes due to increased density (reviewed in (142, 143)), and increased physiological stress from competition and increased number of predators during years with larger returns (e.g., (142)) could all influence the differences observed in immune related genes. Further investigations into the nature of these genes in pink salmon may uncover the environmental factors and selective pressures relevant to the evolutionary history of these pink salmon lineages.

#### Organ development/maintenance

Salmon go through nutritional and behavioural changes that require organ-level alterations and maintenance throughout their life-cycle. This can be observed in developing salmon that transition from plankton to other fish as food sources. In the eye, this transition requires the development of new functionality such as night vision to chase prey. One example of such a transition is the change of UV opsins observed in hatched Pacific salmon to blue later in life by opsin changeover (144, 145).

Variation in vision related genes have previously been observed between sockeye salmon populations (77). In Atlantic salmon, *six6*, a gene related to eye development, daylight vision (146, 147), and fertility (148) was also found to be associated with age at maturity (149, 150) and later with stomach fullness during migration (151). These studies suggest that genetic variation influencing organ development, transition, or maintenance are important components influencing salmonid evolution.

Similar to *six6* in Atlantic salmon, Protein tyrosine phosphatase receptor type J (*PTPRJ*) (152), histidine N-acetlytransferase (*hisat*) (153–160), and microtubule-associated protein 9 (*MAP9* or *ASAP*) (161) all appear to play roles in proper vision. The variation in these genes may represent differences in selective pressure between odd and even-years and could be driven by the different population dynamics observed between odd and even-year populations.

Cystathionine gamma-lyase (*CTH*) may have, among other roles, a function in hearing (162–164), and could have been influenced by similar population dynamics as those suggested for vision-related genes. Multidrug and toxin extrusion protein 2 gene (*SLC47A2*) is not related to a specific organ, though it may have a special role in the blood-brain barrier (165, 166). Instead, it may help in removing toxins, which might accumulate in more dense and stressed populations.

#### Behaviour

Fish display consistent behavioural differences from each other, analogous to human personalities (167). Personality variation in a population may represent adaptive solutions to different environmental pressures (167). In high density populations, such as the odd-year populations, more aggressive behaviours during high-density spawning conditions (40) could result in more offspring, but might waste energy in lower-density conditions. Associations to genes related to behaviour have previously been identified among sockeye salmon populations (77), and under selection between wild and farmed Atlantic salmon (168). In the present study, protein-methionine sulfoxide oxidase mical2b (*mical2b*) (169, 170) and cell division control protein 42 homolog (*CDC42*) (171), both putative genes found in the eigenGWAS analysis between even and odd-year pink salmon, have previously been found to be associated with anxiety/reactiveness and schizophrenia, respectively.

Another gene, we were not able to unambiguously identify as a candidate gene from other nearby genes, on LG15_El08.2-20.1 is worth noting because of its association with behaviour. The gene, 5-hydroxytryptamine receptor 2B (*5-HT2B* and also known as serotonin receptor 2B), is associated with impulsivity and impulsive aggression in humans and mice (172, 173). This region of the genome was fixed in the even-year class pink salmon where population density is often lower (in many North American populations, but not necessarily in all of the rivers in this study) and aggression might be maladaptive. These associations suggest that population dynamics might influence the frequency of certain personality traits in pink salmon populations, with density as a possible driving force.

### Sex determination and sdY

With the discovery of a novel sex-determining gene in salmonids (111), and previously with closely linked genetic markers (174, 175) researchers have been able to identify instances of sex-determination switching between chromosomes in salmonids (176–180). As suggested in Yano et al. (2013), Y-chromosome switching may act in response to (expected) degeneration of the Y-chromosome due to mutation accumulation from reduced recombination (181). In pink salmon, *sdY* was located on LG20_El14.2, but we suggest there may be an alternative location as well.

Several pieces of information indicate that LG20_El14.2 may not be the only location of the sex-determining gene, *sdY*, in the pink salmon genome. For instance, there were two *sdY* haplotypes and several males had multiple copies of this gene. Also, all males with the CGGA *sdY* haplotype had a run of homozygosity similar to most females on the LG20_El14.2 chromosome near the putative location of *sdY*. It is expected that near the *sdY* gene, recombination is reduced and mutations would accumulate between the X and Y-chromosomes as a result of reduced recombination. Females tend to have long runs of homozygous genotypes where recombination is reduced and males tend to have long stretches of heterozygous genotypes when reads from the X and Y-chromosome align at the same location (77). Since the males with the CGGA *sdY* haplotype have long runs of homozygous genotypes at the LG20_El14.2 region, as most of the females do, we suggest that the CGGA *sdY* is at another location in the genome in these individuals. We were unable to identify a precise putative alternative location because there were too few individuals with the CGGA *sdY* to obtain a signal from a genome wide association analysis, however, the potential discovery of another salmon species with alternative *sdY* locations, further supports the hypothesis of Y-chromosome switching put forth by Yano et al. (2013) for salmonids (181).

## Conclusions

We generated reference genome assemblies for both pink salmon lineages, RNA-seq data for genome annotation, and whole genome re-sequencing data to expand the available resources for this commercially important and evolutionarily interesting species. The coupled whole genome re-sequencing study of 61 individuals from several streams in British Columbia (and one from Japan) helped us to characterize regions of the genome that have diverged between the temporally isolated groups. The amount and degree of lineage-specific genomic variation suggests that there is little gene-flow between the year-classes, but the shared variants such as whole mitochondrial and *sdY* haplotypes suggests that there has been enough recent gene-flow or alternative year-class replacement to maintain these similarities. Divergence at centromeres between the two lineages may be a consequence of centromere drive (or genetic drift and reduced recombination) and represent early stages of speciation. Genes related to the immune system, organ development/maintenance, and behaviour were divergent between odd and even-year classes as well. These example lineage defining differences offer us a glimpse into the evolutionary landscape and the selective pressures or demographic histories that may have driven the divergence in these genes and the differences between odd and even lineages of pink salmon.

## Acknowledgements

Extensive sample preparation and sequencing was performed at McGill University and Génome Québec Innovation Centre (now the Centre d’expertise et de services Génome Québec) and we would like to thank the staff and scientists there for their efforts. We would also like to thank the generous computing resources provided by Compute Canada (www.computecanada.ca). Fisheries and Oceans Canada and the University of Victoria facilities and personnel made this work possible. Finally, the authors would like to thank the many Fisheries and Oceans Canada staff who collected samples for analysis in this study.

## Supporting information

**S1 File. Sample information**. The sample tab has metadata about each sample, including information on sex, river, and year-class (latitude and longitude locations are approximate). The StatsAllFilters shows metrics from the .vcf file after filtering for LD (see methods). Stats1stFilter has the same information, but from the .vcf file after only preliminary filtering (see methods). The eigenGWAS tab contains the DAPC values used in the eigenGWAS analysis (see methods). The Mitochondrion tab shows metadata used to generate the mitochondria figures. The GPS tab shows the coordinates used in the sample map. The Admixture tab has the values output from the admixture analysis. For each tab with LG, these sheets have manually genotyped areas and calculations of HWE. The PrivateAlleles tab has metrics output from Stacks. The SharedAlleles tab has a matrix of shared alleles between individuals in long format and statistics on the right. The Y-Chrom tab has information about the *sdY* haplotypes. The GWAS tab has metadata used in the GWAS analysis. The GHp tab displays the alignments of the growth hormone pseudogene and *sdY* gene to the odd and even genome. The even genome placements will change after processing by NCBI.

**S1 Fig. Chromosomal polymorphism at centromere on LG15_El08.2-20.1.** Depiction of LG15_El08.2-20.1 and a chromosomal polymorphism, either a deletion or evidence of a chromosomal fusion. At the top, LG15_El08.2-20.1 is depicted with the distance and location of the purposed polymorphism. Scaffolds/contigs that comprise the region surrounding the polymorphism are shown below the chromosomal depiction, with a blue arrow showing where multiple small contigs were placed. Below the scaffolds, synteny with rainbow trout and Northern pike is shown based on CHROMEISTER (106) alignments. Finally, ONT/Nanopore reads that were used to generate the genome assemblies were aligned back to the odd-year genome and visualized with IGV. Reads in the odd-year individual are shown flanking the deletion (the display was split because the region was too large to adequately visualize otherwise).

**S2 Fig. Sex determining region of the even-year pink salmon compared to the rainbow trout Y-chromosome.** A) A CHROMEISTER (106) dotplot between the Y-specific portion (top) and shared portion (bottom) of LG20_El14.2 of the even-year pink salmon genome assembly and the rainbow trout Y-chromosome (63). The location of the *sdY* gene is shown based on the position in the rainbow trout chromosome and the genes annotated by NCBI are shown on the x-axis at the bottom. B) A plot of the Hi-C contact map of the even-year pink salmon genome assembly produced by Juicebox (64). There are multiple inversions between the pink salmon and rainbow trout genome, but the contact map supports the order and orientation for the pink salmon genome assembly and these could represent actual inversions between species instead of assembly errors.

**S3 Fig. Sex determining region of the even-year pink salmon compared to the coho salmon chromosome 29 autosome.** A) A CHROMEISTER (106) dotplot between the Y-specific portion (top) and shared portion (bottom) of LG20_El14.2 of the even-year pink salmon genome assembly and coho salmon chromosome 29. B) A plot of the Hi-C contact map of the even-year pink salmon genome assembly produced by Juicebox (64). There are multiple inversions between the pink salmon and coho salmon genome, but the contact map supports the order and orientation for the pink salmon genome assembly and these could represent actual inversions between species instead of assembly errors.

**S4 Fig. Sex determining region of the even-year pink salmon with genotype information.** Genotypes are shown from an IGV (107) screenshot for the 61 samples of pink salmon for the region with the *sdY* sex-determining gene. The top portion shows the distance of the Y-specific genome region (∼3.2 Mbp) and the contig/scaffold boundaries that make up this region are shown as vertical lines. Below the distances, allele frequencies for each locus are shown, and below that individual genotypes. The x-axis of the genotypes represent loci and each line on the Y-axis represents an individual pink salmon. The dark-blue colour is a homozygous reference genotype, the light-blue colour a heterozygous genotype, and the green genotype is for a homozygous alternative locus. There are large stretches (1-2 Mbp) of heterozygosity and homozygosity based on sex. Please note that there is a possible inversion (from a mis-assembly) in this region as the runs of homozygosity and heterozygosity are broken by a section from ∼600 kbp and ∼1,300 kbp.

## Notes

### Competing Interest Statement

The authors have declared no competing interest.

